# KegAlign: Optimizing pairwise alignments with diagonal partitioning

**DOI:** 10.1101/2024.09.02.610839

**Authors:** A. Burak Gulhan, Richard Burhans, Robert Harris, Mahmut Kandemir, Maximilian Haeussler, Anton Nekrutenko

## Abstract

Our ability to generate sequencing data and assemble it into high quality complete genomes has rapidly advanced in recent years. These data promise to advance our understanding of organismal biology and answer longstanding evolutionary questions. Multiple genome alignment is a key tool in this quest. It is also the area which is lagging: today we can generate genomes faster than we can construct and update multiple alignments containing them. The bottleneck is in considerable computational time required to generate accurate pairwise alignments between divergent genomes, an unavoidable precursor to multiple alignments. This step is typically performed with lastZ, a very sensitive and yet equally slow tool. Here we describe an optimized GPU-enabled pairwise aligner KegAlign. It incorporates a new parallelization strategy, diagonal partitioning, with the latest features of modern GPUs. With KegAlign a typical human/mouse alignment can be computed in under 6 hours on a machine containing a single NVidia A100 GPU and 80 CPU cores without the need for any pre-partitioning of input sequences: a ∼150× improvement over lastZ. While other pairwise aligners can complete this task in a fraction of that time, none achieves the sensitivity of KegAlign’s main alignment engine, lastZ, and thus may not be suitable for comparing divergent genomes. In addition to providing the source code and a Conda package for KegAlign we also provide a Galaxy workflow that can be readily used by anyone.

## Introduction

International efforts such as the Earth BioGenome Project aim to produce reference genomes for all ∼1.8 million known eukaryotic species over the next decade [1–4]. Such initiatives combined with continuing improvements in sequencing techniques and assembly methods ensure steadily increasing growth in the number of fully sequenced genomes. However, a sequenced genome is just a file with little utility on its own. Comparisons to other genomes is critical for understanding its functional landscape in order to identify constrained (functional) and rapidly changing (evolving) regions. Performing such comparisons in practice means aligning sequences and involves three steps: (1) generation of pairwise local alignments, (2) filtering and ordering of local alignment blocks, and (3) generation of multiple alignment blocks from filtered and ordered local alignments. The first step necessarily involves local alignments, because frequent and ubiquitous structural changes: duplications, inversions, deletions, insertions at multiple scales as well as chromosomal fissions and fusions disrupt the linear order of genomes. Once local alignments are computed, they are post-processed to remove spurious hits, and grouped into chains and nets [5] to define larger syntenic regions. A chain is an ordered sequence of local alignments; and a net is a hierarchical collection of chains covering nucleotides of a given genomic region once [6]. Pairwise alignments representing homology anchors between multiple species are then further combined into multiple sequence alignments (MSA) using different methods (see [7,8]).

**Table 1.** Historic data from 4,780 pairwise alignments generated between 2005 and 2020 by the UCSC Genome Browser team.

| Genome pair count | Genome Sizes | Average CPU Hours |
| --- | --- | --- |
| 1,372 | both large ($> 1$ Gbp) | 2,764.1 |
| 850 | both small ($\leq 1$ Gbp) | 216.4 |
| 168 | mixed size | 1,893.5 |
| 2,390 | all sizes | 1,796.8 |

A number of purpose-built tools were developed in the early-to-mid 2000’s for this task: BLAT [6], CHAOS [7], PatternHunter [8], MUMmer [9], and blastZ [9]. A significantly improved derivative of blastZ—lastZ [10]—now used as the primary local-alignment engine for many production-level multiple alignment pipelines, including multiZ [11] and ProgressiveCactus [12], due to its superb sensitivity across a wide range of sequence divergence levels (see Results). However, lastZ was not built for speed: recent benchmarks put the runtime of a mammalian whole-genome alignment at around 2,700 CPU hours (Table 1). Therefore, reproducing the standard UCSC multiZ alignment of 100 vertebrates using the human genome as the reference would take around ∼30 CPU years. This makes lastZ the main bottleneck preventing MSA generation at scale and complicating our ability to keep up with the rapid pace of multiple large genome sequencing initiatives.

Because of this limitation currently only a few research groups in the world are able to routinely generate whole genome alignments. There are two reasons for that. First, because alignment with lastZ is slow, it is parallelized by dividing input sequences into smaller fragments and running resulting jobs on separate nodes of a large computing cluster. This approach creates logistical challenges stemming from the need to manage hundreds of thousands of distinct alignment runs—a job requiring considerable system engineering and administration expertise. Second, appropriate computational infrastructure is needed to compute alignments. Access to modern clusters remains a hurdle (both financially and logistically) for many research groups even in developed countries, as it requires expertise in resource procurement, configuration, and maintenance along with fiscal resources.

Our goal was to make pairwise genome alignment universally accessible by optimizing it such that it can be performed on a single compute node within several hours. To achieve this goal we initially chose a GPU-accelerated replacement for lastZ called SegAlign [13]. It accelerates the ‘Seed’ and ‘Filter’ stage of the alignment process (for a detailed review of the seed-filter-extend paradigm see [10]). However, after initial tries we discovered that SegAlign does not utilize hardware resources effectively. Specifically, often all hardware resources—both CPU and GPU—wait for a few long-running CPU threads to complete their work. In extreme cases, a single CPU thread can work for over 93% of the total runtime while all other hardware resources—both CPUs and GPUs, stay idle. This problem cannot be fixed by adding more cores or GPUs: doing so exacerbates the problem by resulting in an even larger fraction of idling resources. We also observed that this underutilization is input dependent. Highly similar genomes are more affected by this problem, due to more work done for ‘gapped-alignment’ in the ‘extend’ stage resulting in a CPU bottleneck, rather than GPU.

In this manuscript we refactor SegAlign by introducing diagonal partitioning and leveraging the latest Nvidia GPU features—Multi-Instance GPU (MIG) [25] and Multi-Process Service (MPS) [26]. The new tool, KegAlign, is distributed via a Conda package and our Galaxy system by allowing production-scale alignment generation. This service is freely and readily available to anyone in the world with an Internet connection.

## Results and Discussion

### lastZ is still the most sensitive tool for aligning divergent genomes

lastZ has been around for ∼20 years. In this timeframe new tools such as minimap2 [14] or fastga [15] have been developed that outperform lastZ dramatically in terms of speed (Table 2).

**Table 2.** CPU time required for aligning chromosome 1 from assembly hg38 against chromosome 1 from assemblies hs1 and mm39. Alignment is performed using a single thread on an Intel i9 workstation. A single thread was used to make comparison with lastZ “fair” as it supports single thread only. The time is computed using the time UNIX command. “Real” time is reported here.

| Tool | chr1 hg38 / chr1 hs1 | chr1 hg38 / chr1 mm39 |
| --- | --- | --- |
| lastZ | 867 min 59 sec | 208 min 8 sec |
| minimap2 | 6 min 9 sec | 0 min 36 sec |
| fastga | 1 min 41 sec | 1 min 8 sec |

So why use it? The key advantage of lastZ is its ability to find alignments between highly divergent sequences. To demonstrate this we created a set of “homologous” sequence pairs of varying lengths (1, 2, 5, and 10 thousands nucleotides) with divergence varying from 0% to 40% in 1% increments (164 sequence pairs in total; see Methods). Random 1,000 nucleotide flanks were added to 5’ and 3’ of each sequence. Fig. 1A shows the relationship between alignment coverage (the fraction of the sequence that is included in the alignment) and divergence/length. Only lastZ consistently generates end-to-end alignments across all divergence and length categories. To verify that these lastZ advantages are observed with real data we generated whole genome alignments between hg38 and mm39 builds of human and mouse genomes, respectively. We then computed the fraction of coding exons covered by alignments. In both cases overlapping alignments and exons were merged to eliminate redundancy (see Methods). lastZ alignments covered the absolute majority of protein-coding exons (Fig. 1B) suggesting that it is well suited for comparing genomes from evolutionarily divergent species. Since the genomes of thousands of species sequenced within the framework of Earth BioGenome Project and other initiatives are expected to have the full range of evolutionary distances lastZ remain a critical tool for generation of pairwise alignments.

**Figure 1.**
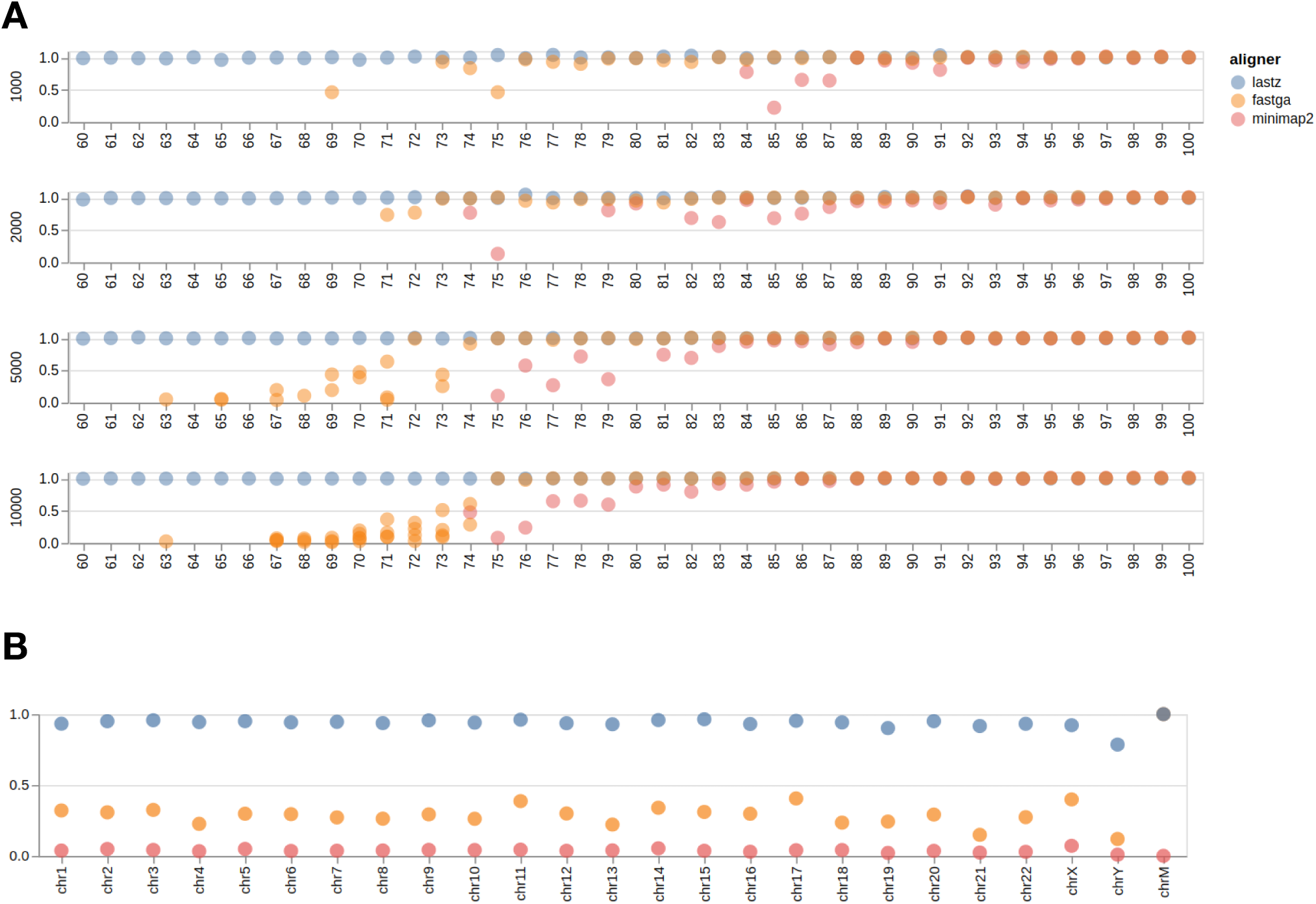
**A**. Comparison of three different pairwise aligners on a set of simulated sequences with different divergences (60 - 100) and lengths (1,000, 2,000, 5,000, and 10,000). The Y-axis (0 - 1) indicates what fraction of the sequence is covered by alignments generated by a given tool. This calculation excludes randomly added flanks. **B**. Fraction of human protein-coding exons (hg38) overlapping with alignments to mouse genome (mm39) produced by each of the three mappers. Coordinates of exons and alignment blocks were merged to avoid redundancy (see Methods).

### GPU-accelerated aligners exist but require optimization

Goenka et al. implemented an accelerated version of lastZ called SegAlign [13]. It uses a graphical processing unit (GPU) to speed up the main lastZ bottleneck (Fig. 2): the Seed-and-Filter stages of the alignment process. The two stages are parallelized on GPU and send their output to the Gapped Extension stage, which is performed by an unchanged lastZ executable. This improves the speed of pairwise alignment by a factor of 5-10✕ [13].

**Figure 2.**
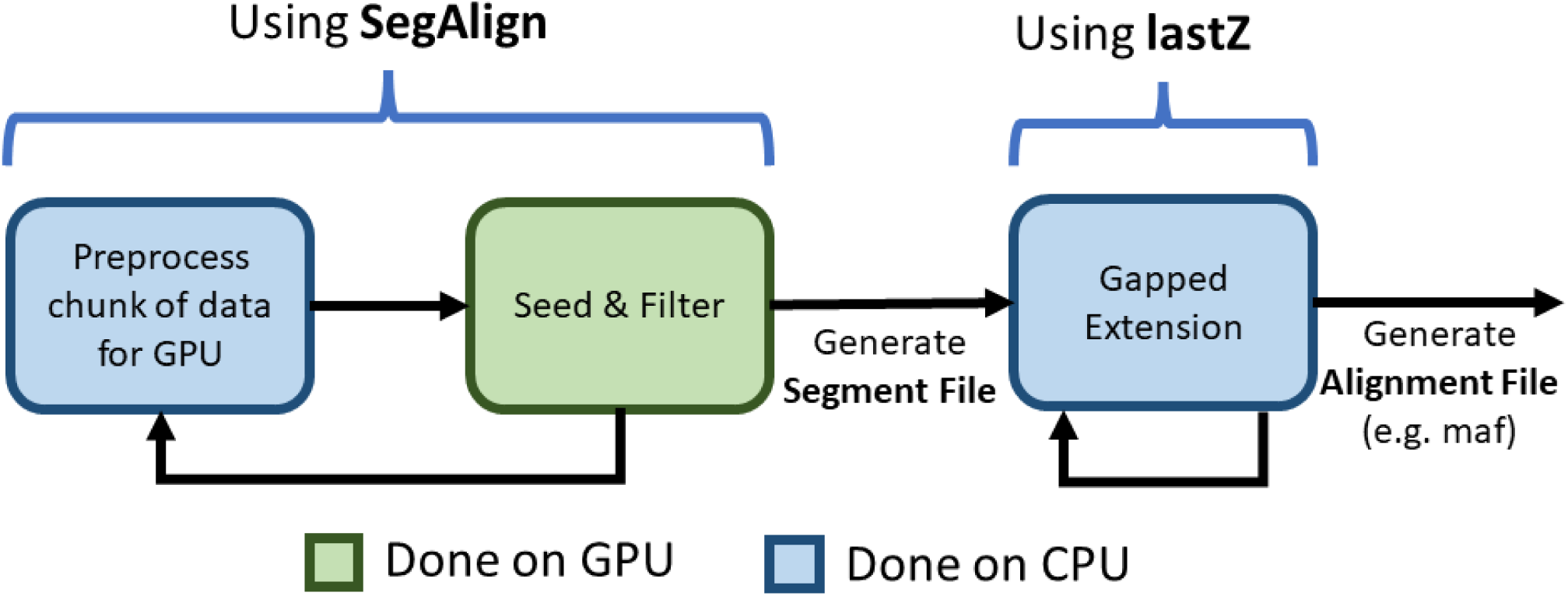
A high-level representation of SegAlign’s pipeline. Blue and green boxes represent steps performed on CPU and GPU respectively. SegAlign repeatedly takes a chunk of the input genomes, performs preprocessing on CPU to make it amenable to run on GPU, and performs the Seed and Extend steps to obtain a segment file, which contains a list of coordinates. Whenever a segment file is output, lastZ is called to perform the Extend stage (i.e. gapped-alignment using the Y-drop algorithm on the coordinates [10]).

As shown in Fig. 2 SegAlign separates the alignment process into GPU- and CPU-bound components. The amount of computation dedicated to each component depends on the sequence divergence between the two genomes being aligned. For inputs with high divergence (e.g. Human vs. Mouse) GPU becomes the bottleneck. This occurs since there are fewer matching alignments for gapped extension. Conversely, for inputs with low divergence (e.g. Human vs. Chimp), CPU becomes the bottleneck due to having many more alignments for gapped extension.

### The CPU underutilization problem

SeqAlign splits input sequences into smaller, equally sized segments and processes them in parallel as separate tasks. However, using equally sized segments does not necessarily lead to equal amounts of work to be performed on each segment pair. The difference in work can be observed in the Gapped-Extension part of the alignment, which is caused by the final number of alignments in a given region of both genomes. Imbalanced partitions can lead to significant under-utilization of hardware resources due to what we call tail latency. For example, in the case of human vs. chimpanzee alignments (Fig. 3) CPU is idle over 50% of the total run time. In fact, for pairs of genomes with low divergence such as human/chimp some regions can have large numbers of seeds, which can lead to a disproportionately large amount of time spent aligning these regions (10,000 times longer than the median runtime in some cases; Fig. 4; also see Supp. Fig. 1). Fig. 4 illustrates the disparity of runtimes in different regions for a human-chimp alignment. This results in the entire alignment not completing until the most time-consuming pair of chunks are completed, which is a typical example of the tail latency problem. Adding additional resources does not help with this problem, since each pair of regions is assigned to a single CPU, and additional CPUs stay idle waiting for these pairs of chunks to finish (Fig. 3).

**Figure 3.**
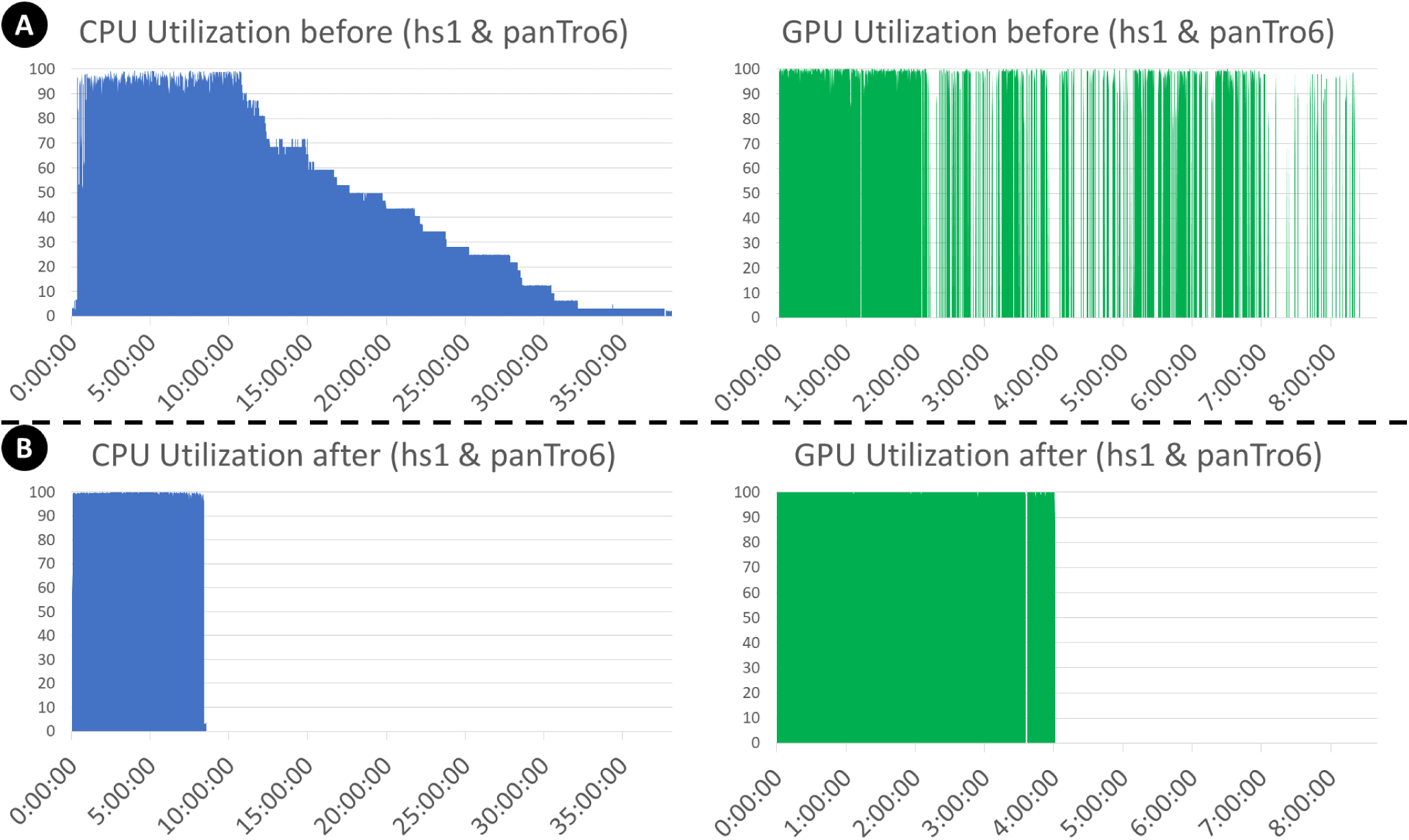
The CPU and GPU utilizations of 32 CPUs and 1 A100 GPU for aligning human (hs1) and chimpanzee (panTro6) genomes. **A**. Severe CPU- and GPU-underutilization of the original SegALign implementation. This alignment completes in 38 hours. **B**. A SegAlign implementation modified by our group allows the same alignment to finish in 8.5 hours.

In fact, for pairs of genomes with low divergence such as human/chimp some regions can have large numbers of seeds, which can lead to a disproportionately large amount of time spent aligning these regions (10,000 times longer than the median runtime in some cases; Fig. 4). This results in the entire alignment not completing until the most time-consuming pair of chunks are completed, which is a typical example of the tail latency problem. Adding additional resources does not help with this problem, since each pair of regions is assigned to a single CPU, and additional CPUs stay idle waiting for these pairs of segments to finish.

**Figure 4.**
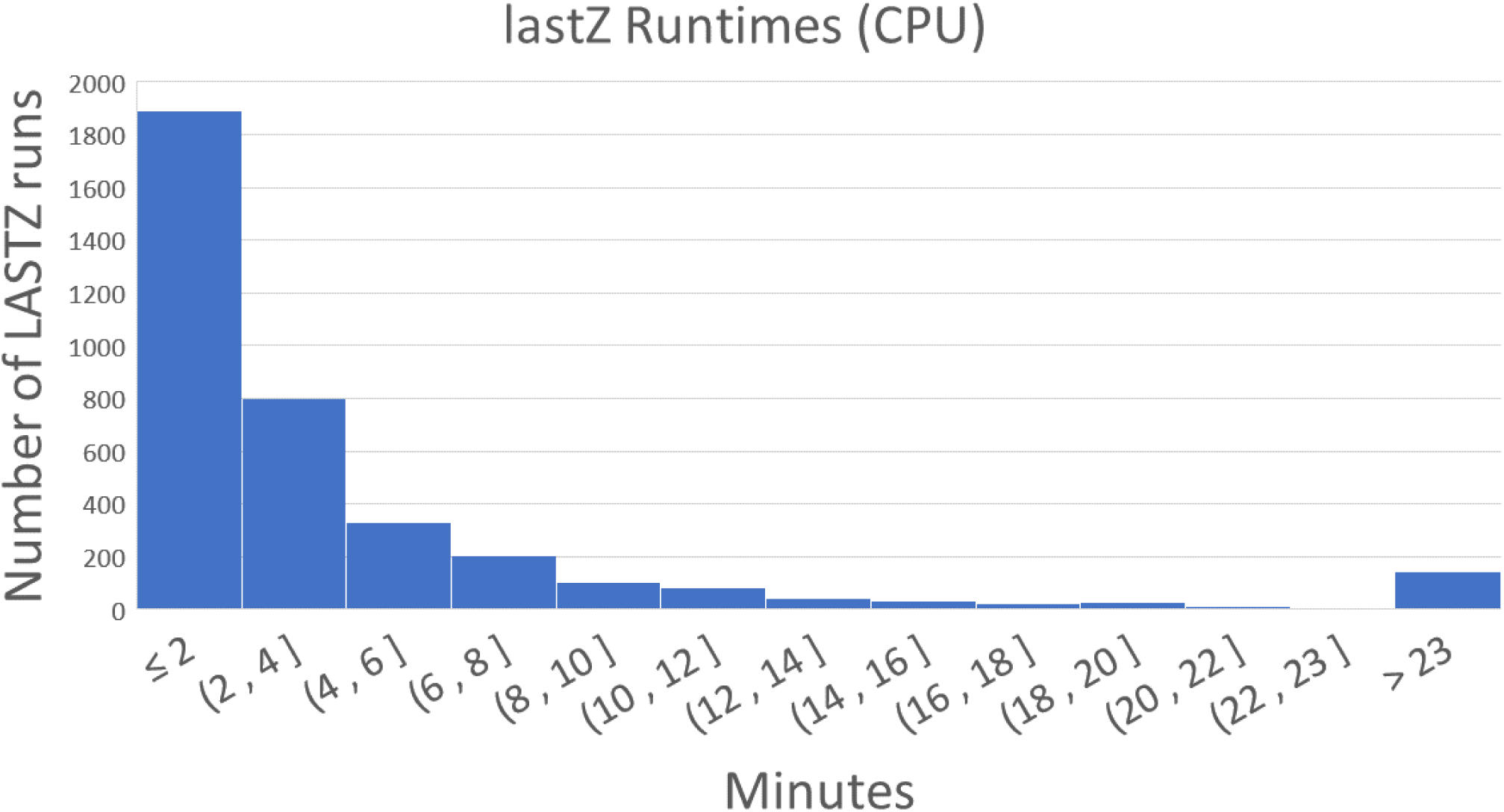
lastZ runtimes for hs1 (human) and pantro6 (chimp). The longest runtime is 27 hours.

### The GPU underutilization problem

We observed two different cases where GPU is underutilized. For inputs with low divergence, there is not enough incoming work to fully utilize the GPU. The first case can be observed in a human-chimp alignment in Fig. 3A where GPU utilization sharply decreases after ∼2 hours. This case is due to GPU data starvation—not enough incoming work to fully utilize GPU—and is caused by two factors. First, CPU starvation due to a large amount of CPU work in the gapped extend stage monopolizing all of the CPU compute, delaying the preprocessing required to begin work on GPU (Pre-process stage in Fig. 2). Second, communication deadlock is caused by the inter-process communication buffer (UNIX pipe) running out of space, due to work arriving much faster than CPUs can execute them. This leads to lastZ output commands (written to the stdout buffer) in SegAlign being blocked, preventing additional GPU work from beginning, as the SegAlign process is waiting for free space in the communication buffer before continuing. These not only lead to an increased runtime, they also cause the GPU to idle for long periods of time, up to several hours, preventing other users and applications (potentially other alignments) from utilizing the GPU. For the second case of GPU underutilization, we observe that GPU utilization ranges between 90 to 100% for all types of inputs even without GPU starvation, as seen in Fig. 3B in the first ∼2 hour period. This is caused by small inefficiencies in the GPU kernel (code) used in SegAlign leading to decreased GPU utilization. While this can be solved by rewriting the GPU kernels more efficiently, we instead chose to take advantage of new NVIDIA GPU features MIG and MPS to increase GPU utilization without changing SegAlign’s code [25, 26].

**Figure 5.**
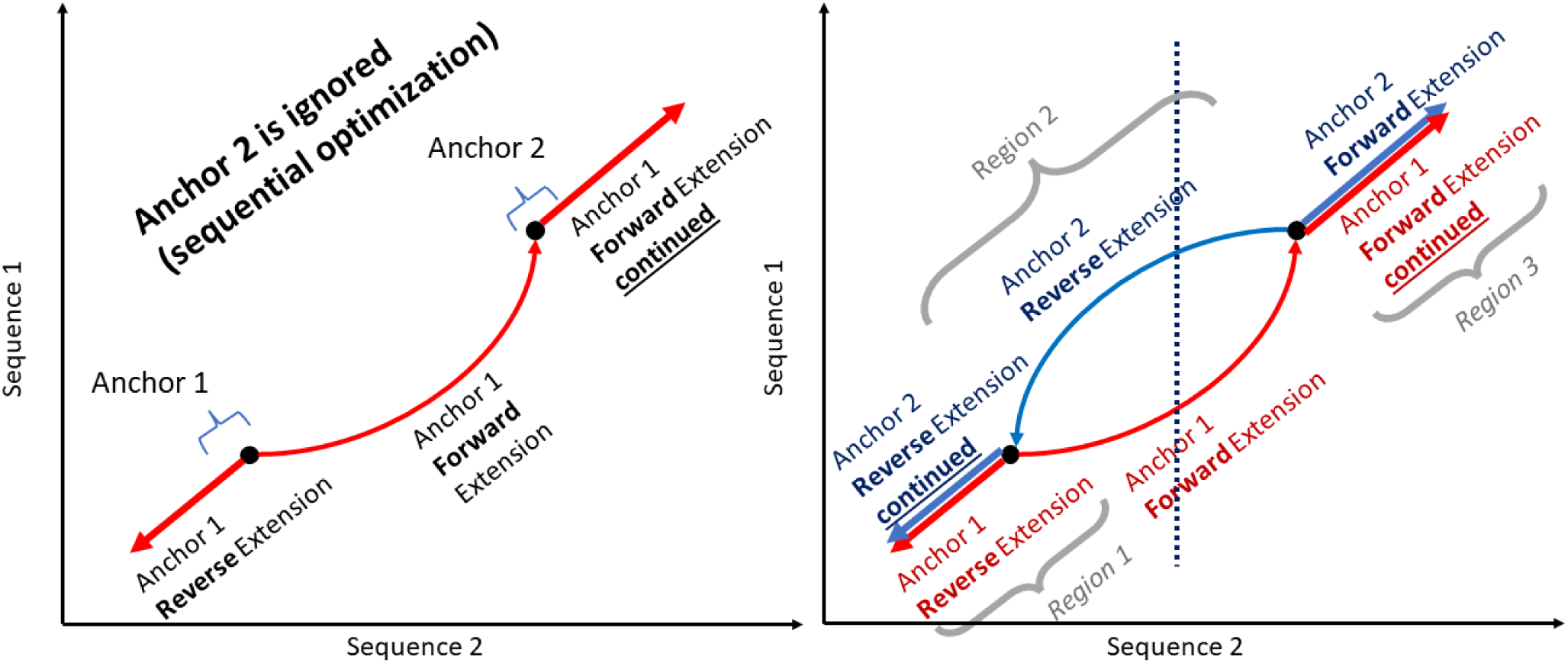
Example showing how straddling alignments can form. Each figure represents a dot plot which is commonly used to depict WGA alignments. Fig. (a) shows the behavior of two anchors which are located on a positive alignment (direction ‘+’ in their segment file rows). Anchor 1 is extended in both directions and crosses over anchor 2, which as a result is not extended. Fig. (b) shows the two dependent anchors located in separate partitions (i.e. segment files) as a result of a column-wise partitioning strategy, depicted by the dotted line. Each anchor is processed independently in parallel, thus both anchors have no knowledge of the other. This leads to two separate extensions, denoted by the red and blue lines. Extensions with the same direction across the same region (regions 1 and 3) result in overlapping alignments. Extensions with different directions across the same region (region 2) result in additional alignments — both alignments follow two equally scoring paths, which are caused by different tiebreakers in the dynamic programming matrix used to calculate them.

### Solving the CPU underutilization challenge with diagonal partitioning

SegAlign, which runs on GPU, performs the seed and extend steps of the alignment process (Fig. 2) and outputs segment files which contain High Scoring Ungapped Alignment Pairs (HSPs). In SegAlign each segment file contains all of the HSPs from a square chunk in the alignment matrix, which by default is 10✕10 MB (set using the lastz_interval_size parameter). A lastZ process is run for each segment file generated by SegAlign and it runs on a single CPU. However, the number of HSPs within this chunk can range from one to over 10 million and thus can cause dramatic variability in runtimes, as illustrated in Supp. Fig. 1. This, in turn, leads to a tail latency problem in which GPUs and most CPUs cores stay idle while few CPUs (running lastZ threads) remain occupied. As can be seen from Fig. 4 nearly all lastZ executions are completed in less than 10 minutes, however the longest execution takes over 27 hours. Thus the entire runtime is lower bound by the longest execution time (Fig. 3).

To solve the tail latency problem, we limited the maximum work for a given lastZ run by restricting the number of HPSs in the segment File. If the number of HSPs is above a certain specified threshold, the segment file is split into two and so on. However, naively splitting segments (blocking, row-wise partitioning, etc.) leads to overlapping alignments and increases output file size (up to several times larger). This is illustrated in Fig. 5. One issue is that lastZ keeps track of previous extended anchors—a point residing on an HSP where ungapped alignment begins—in the input segment file. If an extension goes through another anchor (which is chosen from the segment), that anchor is not extended later. If these segments/anchors are in separate files (and thus will be processed by separate lastZ runs), then both anchors are extended, leading to overlapping alignments. Furthermore, in lastZ, the dynamic programming tiebreakers for a forward and reverse extension are different (each anchor is extended both forward and backward), so the additional segments/anchors being extended can also lead to additional alignments due to different (but equally scoring) paths being taken. It is best to partition in a way that anchors which would be located on the same alignment, are also located in the same segment file. Thus, a fundamentally different approach to partitioning is necessary.

We observe that dependent anchors occur across diagonal regions, since alignments are extended diagonally overlapping, thus dependent anchors occur only along the alignment direction. Anchors are extended in a forward diagonal, for forward alignments, and a reverse diagonal, for reverse alignments, with few horizontal and vertical movements corresponding to gaps. Therefore, we can choose our partition shape to follow along this diagonal to minimize dependent alignments occurring in different partitions leading to straddling alignments (additional and duplicate alignments which are caused by violating dependencies when partitioning). This results in a partition as shown in Fig. 6. Here each colored region indicates a partition, with an equal number of anchors in each. Ideally we want to have the same amount of computation for each partition, which would result in each partition taking roughly the same amount of time. We empirically observe that, in practice, the number of segments is directly proportional to the time it takes for a partition to complete, thus we use the number of segments as a heuristic to determine the amount of work per partition.

Additionally, each row in a segment file does not contain the exact coordinate of the respective anchors, it contains the start and end positions for both the query and reference genomes which indicate the region of likely homology, but there is the possibility that the optimal alignment may include only some regions of the HSP [24]. Therefore, to take into account this possibility, the HSP is reduced to a single point between these two coordinates called the anchor in which the ungapped alignment will begin.

**Figure 6.**
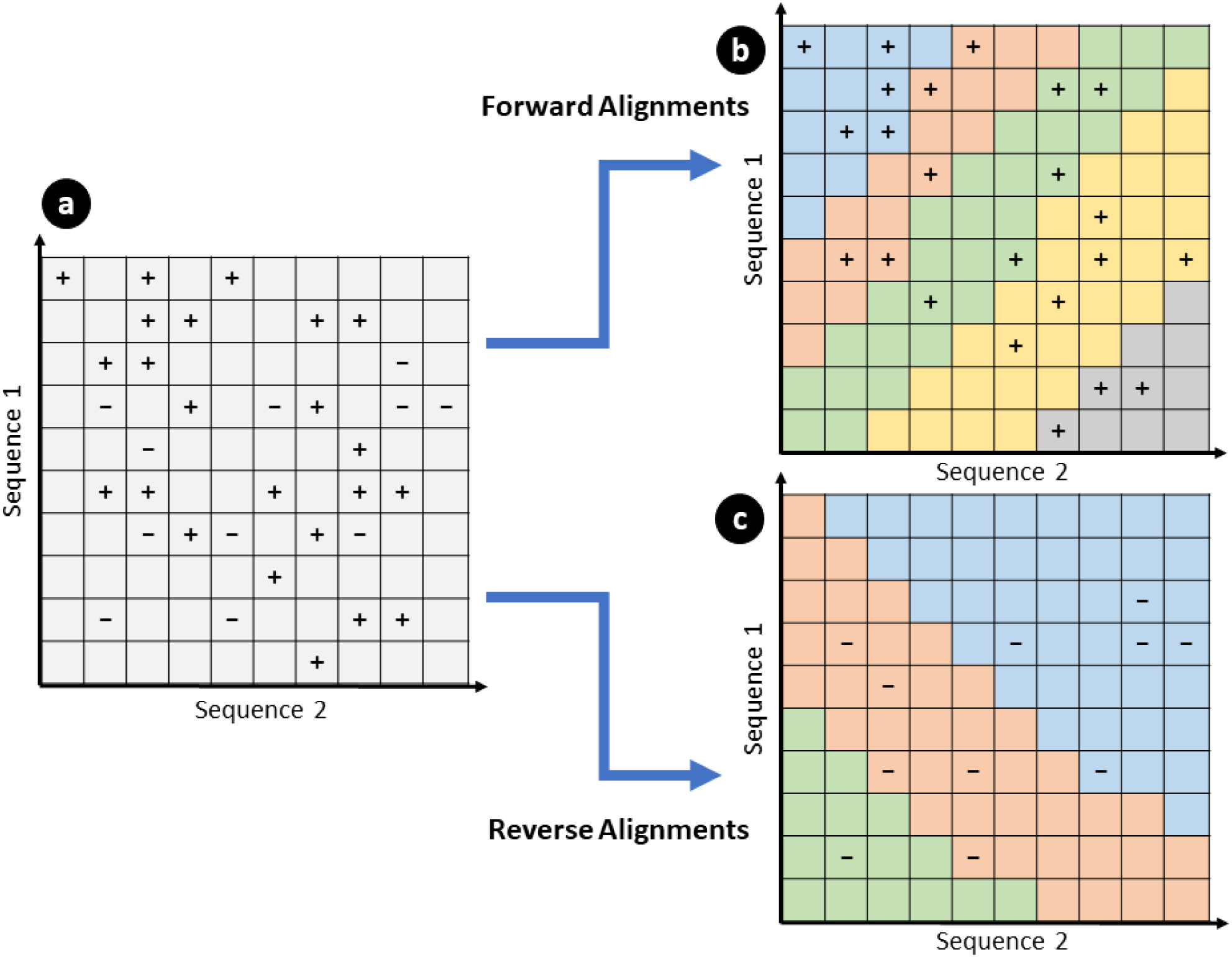
Example of diagonal partitioning for forward and reverse alignments. In (**a**) two genomes of length 10 base pairs are shown, with anchor locations shown as “+” and “-”. “+” represents forward anchors, which will be extended along the positive diagonal and “-” represents reverse anchors, which will be extended along the negative diagonal. (**b**) and (**c**) show separate partitions of a maximum of 5 anchors (also referred to as segments), represented as different colors, for the segments in (**a**), resulting in a total of 8 partitions, each of which will be written into a separate segment file. Since partitions are aligned with the direction of extension, the chances of straddling alignments are minimized. However, it is still possible to have straddling alignments in regions where many anchors are close together, such as in (**b**) where the blue partition ends and the red partition begins.

This is performed in lastZ by calculating the point, along each HSP in the segment files, where the optimal alignment is most likely to include. However, since we do not want to introduce additional overhead when partitioning by calculating anchor positions, we assume that the anchor is chosen in the exact midpoint of the two coordinates and partition accordingly. While our diagonal partitioning scheme does not completely eliminate straddling alignments, it reduces them to a negligible number, as shown below. These mainly occur in regions of the input genomes where many anchors are located within a small region resulting in narrow diagonal segment widths. This, in turn, results in a higher chance of alignments passing through boundaries due to vertical and horizontal movement caused by gaps. SegAlign’s partitioning scheme for Seed and Extend steps also introduces a small amount of straddling alignments which are part of the reported additional alignments in SegAlign’s paper [13].

Diagonal partitioning has an additional benefit of reducing the total amount of CPU work performed in lastZ, as shown in Fig. 7, which further reduces runtime. This is due to lastZ managing a data structure with all ungapped portions of alignments found so far, which is queried during extension of a current anchor to determine bounds. However, operations on this data structure have quadratic time complexity, thus by limiting the number of HSPs in a segment file, the more efficient it is to make operations on this data structure.

Diagonal partitioning introduces a new parameter, the maximum number of segments to include in each partition. The optimal choice to obtain minimum runtime for this parameter depends on two input genomes—their size and similarity—and on the number of CPU cores available. The larger the input genomes and the more similar they are, a larger maximum segment file size is optimal (around 15,000 for human-primates; see Fig. 7). Determining the optimal value for this parameter, for minimum runtime, can be difficult without experimentation. Therefore, we introduce a method to dynamically choose this parameter during runtime, which does not require explicit communication between partitions. Whenever a segment file is made, our partitioning scheme chooses the maximum segment file size as the upper quartile of the original sizes (i.e. before partitioning) of previously generated segments files. Each segment file is written to a shared folder, thus explicitly keeping track of segment sizes is not required. We empirically found that this gives a value within 10% error margin of the optimal maximum segment file size parameter. Additionally, to get the number of lines in a segment file, we divide the file size of each segment file by the size of a single line from the segment file we are partitioning. This gives a close approximation of line sizes without requiring reading through the entire segment file.

### Solving GPU underutilization problem

Above we listed three issues with GPU utilization: CPU starvation, communication deadlock, and inability to fully utilize all available GPU resources. To address the CPU starvation problem we decreased the priority of the lastZ threads (using the Linux nice command) so that the CPU preprocessing work required for the GPU work to begin will not be delayed by multiple threads running gapped-extension. To solve the second problem, communication deadlock, we first identified the root cause – the UNIX pipe buffers used for communicating between SegAlign and lastZ running out of memory. We simply increased the maximum buffer size to solve this problem. For the last problem, inability to fully utilize all available GPU resources, we use Nvidia’s MIG and MPS features [25, 26] to run multiple SegAlign instances in parallel on the same GPU. This results in at least one SegAlign instance fully utilizing GPU resources whenever other instances do not. To accomplish this, we split the input genome across pairs of chromosomes and use the Longest-processing-time-first Algorithm [16] to assign multiple chromosomes into similar sized bins and run a SegAlign instance on each pair of bins. For example, whole genomes contain many chromosomes of different sizes and we assign subsets of these into a bin so that each bin contains around 200 million base pairs (depending on the size of the genome the number of bins may vary). Splitting across chromosomes rather than splitting individual chromosomes in half gives two benefits. First, it prevents additional straddling alignments from occurring, since alignments do not extend across multiple chromosomes. Second, it does not require recalculating alignment coordinates after alignment, which would occur had we split individual chromosomes. We tried different combinations of MIG instance sizes, MPS processes per MIG instances, and partition sizes. We found that having seven MIG instances and up to four MPS instances gives the best results in runtime for smaller sized inputs. For large genome inputs, three or less instances are used if GPU memory is not sufficient. Care must be taken so that GPU memory does not run out when all possible SegAlign instances are running on the GPU. This can be done by limiting the total number of SegAlign instances running on a GPU and by limiting the sizes of the bins for each genome. We used a bin size of around 200 million base pairs, which results in roughly 12 GB of GPU memory used by each SegAlign instance. While this method has the downside of using more GPU memory, devices which support MIG are datacenter GPUs which typically have over 80GB of GPU memory, most of which goes unused when running a single SegAlign instance. For cases where MIG is unavailable, we found using only MPS up to four instances also gives similar speedup.

### Evaluation across evolutionary distances

Fig. 8 shows the runtime results for aligning multiple vertebrate genomes using SegAlign with default parameters and SegAlign with diagonal partitioning optimization. All results were obtained running on 32 Intel Xeon Gold 6330 CPUs and 1 Nvidia A100 GPU, with 200 GB of memory. The results show that genomes with low phylogenetic distance, such as human (hg38 and hs1), aligned with chimpanzee (panTro6), and gorilla (gorGor6) genomes have the greatest improvement from our diagonal partitioning strategy. However, for genomes with high phylogenetic distance, such as human aligned with mouse (mm10 and mm39) and chicken (galGal6) there is little to no improvement using diagonal partitioning. This is because the amount of work for the filter step, for determining the final alignments, depends directly on how similar the aligned genomes are. For the MIG/MPS results, we see that low and high phylogenetic distance genomes benefit from this optimization, but to a much lesser extent than diagonal partitioning; we observe between 10 to 20% improvement in all genomes. However this optimization is complementary to our diagonal partitioning approach and we obtain the best results when combining both optimization strategies.

**Figure 7.**
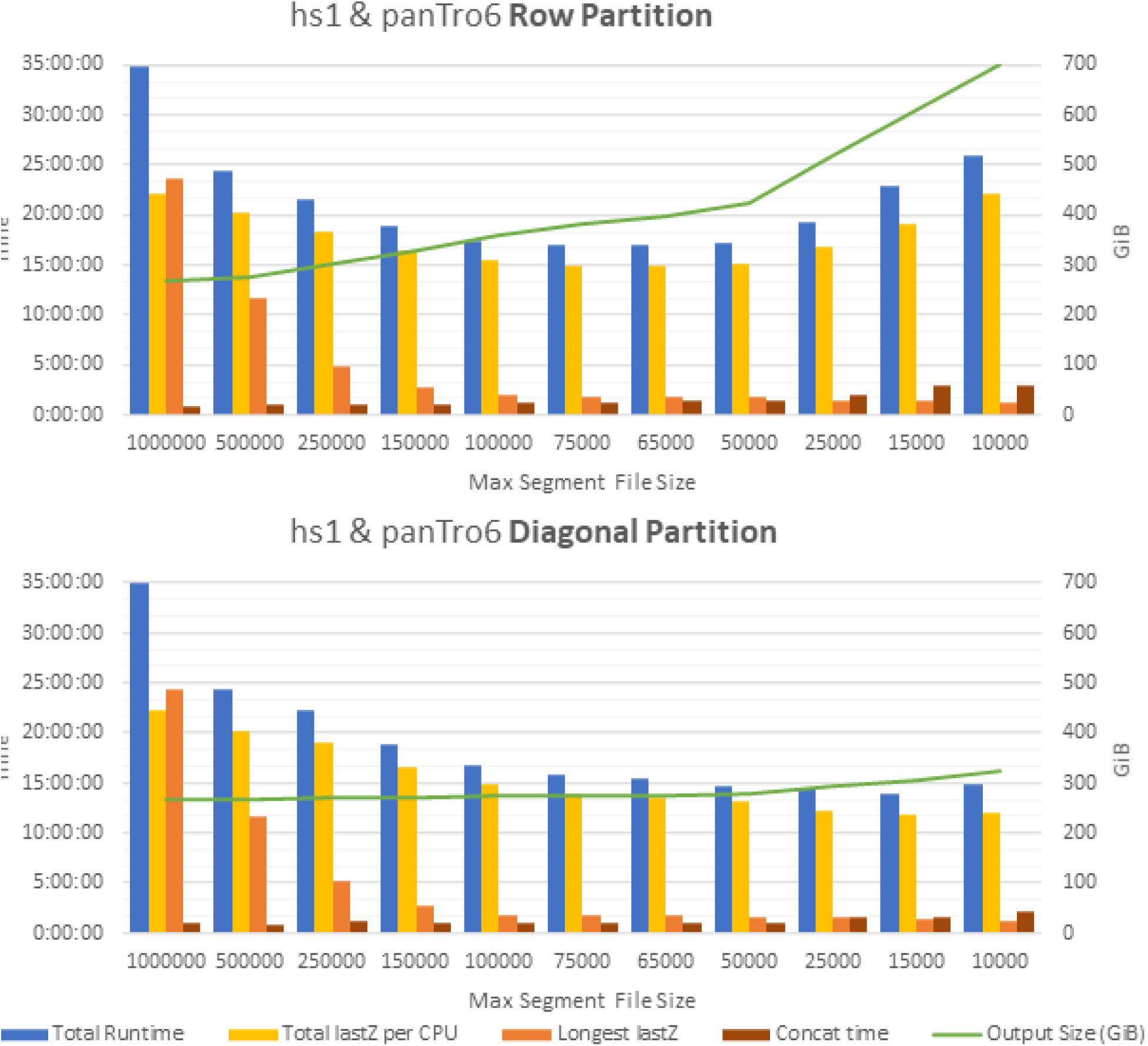
Parameter tuning results for row-wise and diagonal partitioning schemes. Results obtained using 1 GPU and 32 CPUs. Output file size rapidly increases when using row partitioning as maximum segment file size is decreased. Decrease in runtime is due to two factors: (i) decrease in the longest lastZ execution time (Longest lastZ), and the decrease in the total amount of CPU work performed (Total lastZ per CPU). Increased output size leads to slow down from overlaps leading to extra work in lastZ and increased time to concatenate all lastZ outputs into a single file. Baseline values for runtime is 36.5 hours, max lastZ is 27 hours, and output size is 267 GB.

**Figure 8.**
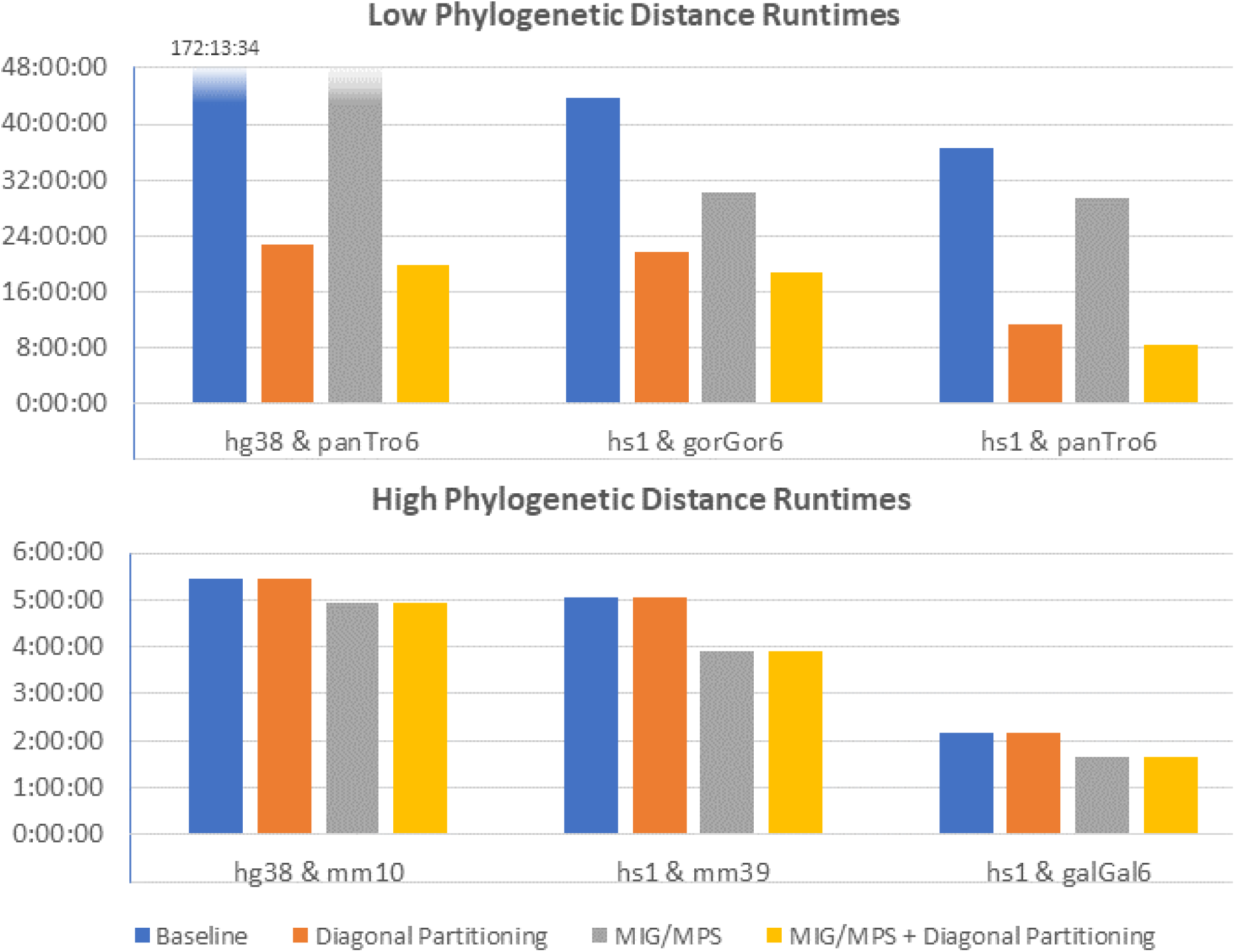
Results for multiple whole genome vertebrate inputs. Genomes with low phylogenetic distance (high similarity) have a large decrease in runtime using segment splitting, while genomes with high phylogenetic distance have little to no improvements. MIG/MPS gives a moderate improvement in all cases. The best results are obtained by combining the two methods.

### Democratizing of genome alignments with KegAlign workflow

To make our work useful to the broad research community we incorporated KegAlign in the Galaxy system developed by our group [17]. It consists of two components: a KegAlign tool and a specialized version of lastZ called “batched-lastZ”. The KegAlign tool performs the “Seed-and-Filter” stage of the alignment generation and is configured (see Methods) to run on a GPU-containing node. The tool outputs a “keg”—compressed tar archive containing partition files with coordinates of segments for performing gapped-extensions and a list of lastZ commands to process these segments. The number of commands and the number of segments depends on size of the input genomes and their divergence. For example, aligning human genome build hg38 against build hs1 generates 5,490 partitions containing a total of 18,227,861 segments. For each of the 5,490 partitions there is a corresponding lastZ command. The batched-lastZ accepts the Keg archive as an input and executes all lastZ commands (in the above example that is 5,490 lastZ commands) listed there. In Galaxy the two tools are combined into a workflow shown in Supp. Fig. 2. The workflow accepts the following inputs: a target sequence, one or more query sequences, and an optional substitution scoring matrix (by default the HOXD70 scores are used [18]). It outputs alignments in a variety of formats that can be specified by the user including MAF (default), BAM, and a variety of configurable tab-delimited flavors. Performance of the workflow’s two main components, KegAlign and batched-lastZ, are shown in Table 3. The tools are run on ACCESS-CI infrastructure as described in *Methods* and are accessible to any user of https://usegalaxy.org instance.

In addition of running tools via the graphical user interface (GUI) they can be run programmatically using Galaxy’s planemo framework [19] against an existing public Galaxy instance as follows:

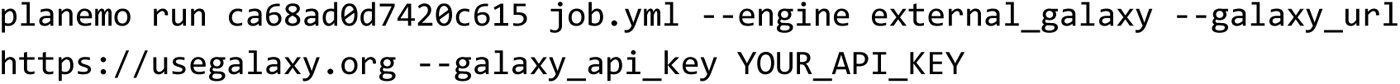

In this case workflow id (ca68ad0d7420c615) corresponds to the KegAlign workflow shown in Supp. Fig. 2. Job.yml specifies inputs and “YOUR_API_KEY” corresponds to a user’s API key on https://usegalaxy.org instance. The key advantage of this model is that not only workflow with all necessary tools is provided, but the infrastructure suitable for computing alignments is provided as well!

The source code of KegAlign can be obtained from https://github.com/galaxyproject/KegAlign or from CondaForge at https://anaconda.org/conda-forge/kegalign.

**Table 3.** Performance of KegAlign/Batched-lastZ pipeline. hg38 & hs1 = *Homo sapiens*; panTro6 = *Pan troglodytes*; mm39 = *Mus musculu*s; galGal6 = *Gallus gallus*; Pf^3D7^ = *P. falciparum* 3D7 (GCA_000002765.3); Py^17XNL^ = *P. yoelii* 17XNL (GCA_020844765.3); Pc = *P. chabaudi* (GCA_900002335.2); Py^17X^ = *P. yoelii* 17X (GCA_900002385.2); Pf = *P. falciparum* (GCA_900632045.1); Pvv = *P. vinckei vinckei* (GCA_900681995.1); Pvb = *P. vinckei brucechwatti* (GCA_903994205.1); Pvp = *P. vinckei petteri* (GCA_903994235.1)

| Target | Query | # alignment blocks | KegAlign (32 cores, 1 A100 GPU) |  | Batched lastZ (80 cores) |  |
| --- | --- | --- | --- | --- | --- | --- |
|  |  |  | Walltime | Peak Memory | Walltime | Peak Memory |
| hg38 | hs1 | 104,308,265 | 5 h 55 m | 26.4 Gb | 6 h 47 m | 134.1 Gb |
|  | panTro6 | 119,368,742 | 5 h 59 m | 28.0 Gb | 9 h 4 m | 124.2 Gb |
|  | mm39 | 10,963,863 | 5 h 16 m | 18.5 Gb | 21 m | 97.7 Gb |
|  | galGal6 | 686,203 | 2 h 15 m | 14.3 Gb | 3 m | 86.1 Gb |
| Pf<sup>3D7</sup> | Py<sup>17XNL</sup> | 40,623 | 60 s | 695 Mb | 6 m | 2.4 Gb |
|  | Pc | 15,671 | 15 s | 586 Mb | 3 m | 1.9 Gb |
|  | Py<sup>17X</sup> | 40,321 | 23 s | 595 Mb | 6 m | 2.4 Gb |
|  | Pf | 131,787 | 16 s | 596 Mb | 29 m | 5.7 Gb |
|  | Pvv | 14,882 | 13 s | 600 Mb | 3 m | 1.8 Gb |
|  | Pvb | 16,827 | 12 s | 585 Mb | 3 m | 1.9 Gb |
|  | Pvp | 20,095 | 60 s | 679 Mb | 4 m | 1.9 Gb |

## Discussion

To our knowledge, there is no prior work on partitioning segment files. There are several works that take into account the diagonal extension property of genome alignment. These methods focus on decreasing the search space of dynamic programming, but none are used for partitioning data. Several papers [20,21] use a method of solving dynamic programming tables called the wavefront algorithm, which takes advantage of the fact that *k*-mers (or seeds) are extended in a diagonal direction. Wu et al. [22] introduces a method called diagonalization, which also takes advantage of the diagonal extension property of *k*-mers and uses this for limiting the search space for “oligomer chaining”, a method for approximate alignment.

The advantage of our approach is in that it offers a *practical* solution to pairwise alignment of divergent genomes that can be readily used via a programmatic or GUI route. In particular, our next task will be developing a Galaxy-based service that would allow submission of pairwise alignments to be performed on our infrastructure using KegAlign workflows *en masse*. It will then be possible to chain the alignments and to proceed to generation of multiple alignments using either ProgressiveCactus or multiZ tools relieving the current bottleneck and enabling multiple genome sequencing efforts to rapidly generate large multispecies alignments. This service will not be limited to KegAlign and will also offer minimap2, fastga, and other pairwise aligners.

## Acknowledgements

We thank Dr. Sneha D. Goenka for explaining implementation details of SegAlign. We also thank Dr. Santhosh Girirajan for providing us with access to a powerful GPU-containing workstation. Nate Coroar and Marten Cech were instrumental in deploying the tools developed here on the Galaxy Platform. Hiram Clawson provided valuable information on generation of pairwise alignments at UCSC. This work is funded by NIH Grants U41 HG006620 as well as NSF DBI award 2419522. We also would like to express our immense gratitude to Dan Stanzione and David Hancock for essential computational resources provided by the Advanced Cyberinfrastructure Coordination Ecosystem (ACCESS-CI), Texas Advanced Computing Center, and the JetStream2 scientific cloud.

## Methods

### Simulating sequences across various divergence levels

We generated random sequence pairs with 1,000, 2,000, 5,000 and 10,000 nucleotides (nt) in lengths and divergences between 0% and 40% in 1% increments. Short indels (ranging from 1 to 5 nucleotides) were introduced at a rate of 1%. Random 1,000 nucleotide flanks were added to each sequence. Scripts used for generation of these sequences can be found at https://github.com/rsharris/echydna. The sequences were aligned using the three aligners using versions and parameters described in Table 4 below:

**Table 4.**
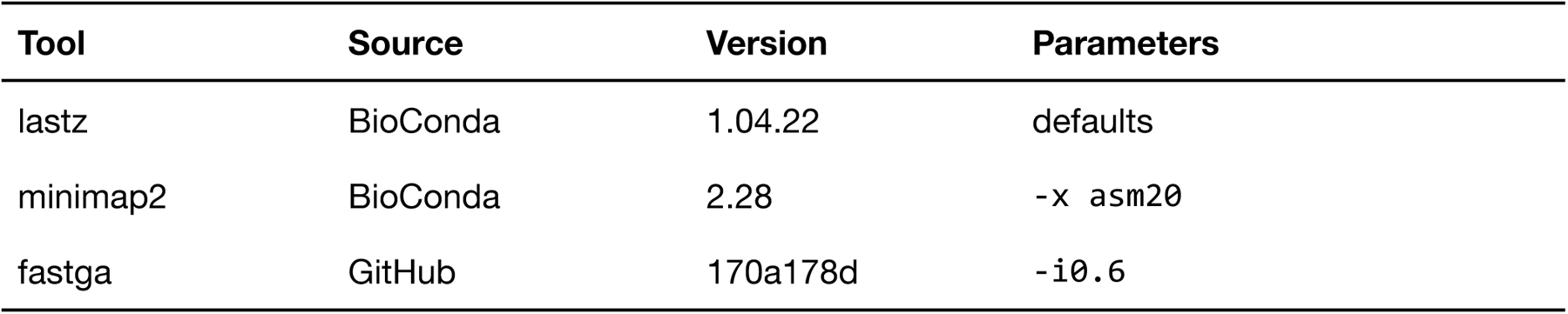
Versions and setting for the three aligners using for testing.

### Coding exon coverage analysis

For this analysis we compared the hg38 build of the human genome against mm39. hg38 was used because it still has the highest annotation density Genome files were downloaded from UCSC:

- hg38 = https://hgdownload.soe.ucsc.edu/goldenPath/hg38/bigZips/hg38.fa.gz
- mm39 = https://hgdownload.soe.ucsc.edu/goldenPath/mm39/bigZips/mm39.fa.gz

We used the same version of minimap2 as specified Table 4. Instead of lastZ we used KegAlign release v0.1.2.7. For fastga we used code repository version ‘74d5666’. Parameters were the same as in Table 4. Resulting datasets were uploaded and analyzed in the following Galaxy history: https://usegalaxy.org/u/cartman/h/aligners--exons.

Postprocessing was performed using https://usegalaxy.org/u/cartman/w/cdsoverlaps workflow. Briefly, the workflow performs the following steps:

1. Re-format files to generate a 3-column bed file containing only coordinates of alignment blocks
2. The intervals in these files are merged using bedtools merge [23] to remove redundancy. The final result is the list of non-overlapping unique intervals for each aligner.
3. A bed file containing coordinates of all protein-coding exons for hg38 (5’- and 3’-UTRs are stripped) is fetched from the UCSC Table Browser and also merged using bedtools merge.
4. Coordinates of alignments from each tool are then projected to coordinates of exons using ‘bedtools annotatè.
5. These data were then aggregated by chromosome by counting the number of exons that overlap alignment blocks for each aligner. The total number on each chromosome was divided by the number of exons covered by alignments obtaining the final fraction for each tool.
6. This data is processed using a Jupyter Notebook to generate Fig. 1B.

All notebooks and other artifacts can be found at https://github.com/nekrut/lz-fg-mm

### Galaxy configuration with TPV

KegAlign and batched-lastZ are configured to run on large compute nodes to achieve good performance. At the time of writing, they were using the following resources:

- KegAlign - g3.medium node at https://jetstream-cloud.org/. The node provides 8 CPUs and 1 A100 GPU as well as 30 and 10 Gb of RAM for CPUs and GPU, respectively.
- Batched-lastZ - ICX “Ice Lake” compute node on Stampede3 cluster at the Texas Advanced Computing Center (TACC) with 40 CPUs (80 cores) and 256Gb RAM.

Jobs executed by Galaxy Tools are routed by the Total Perspective Vortex (TPV) framework (https://github.com/galaxyproject/total-perspective-vortex) and configured as specified in https://github.com/galaxyproject/usegalaxy-playbook.

## Supplemental Figures

**Supplemental Figure 1.**
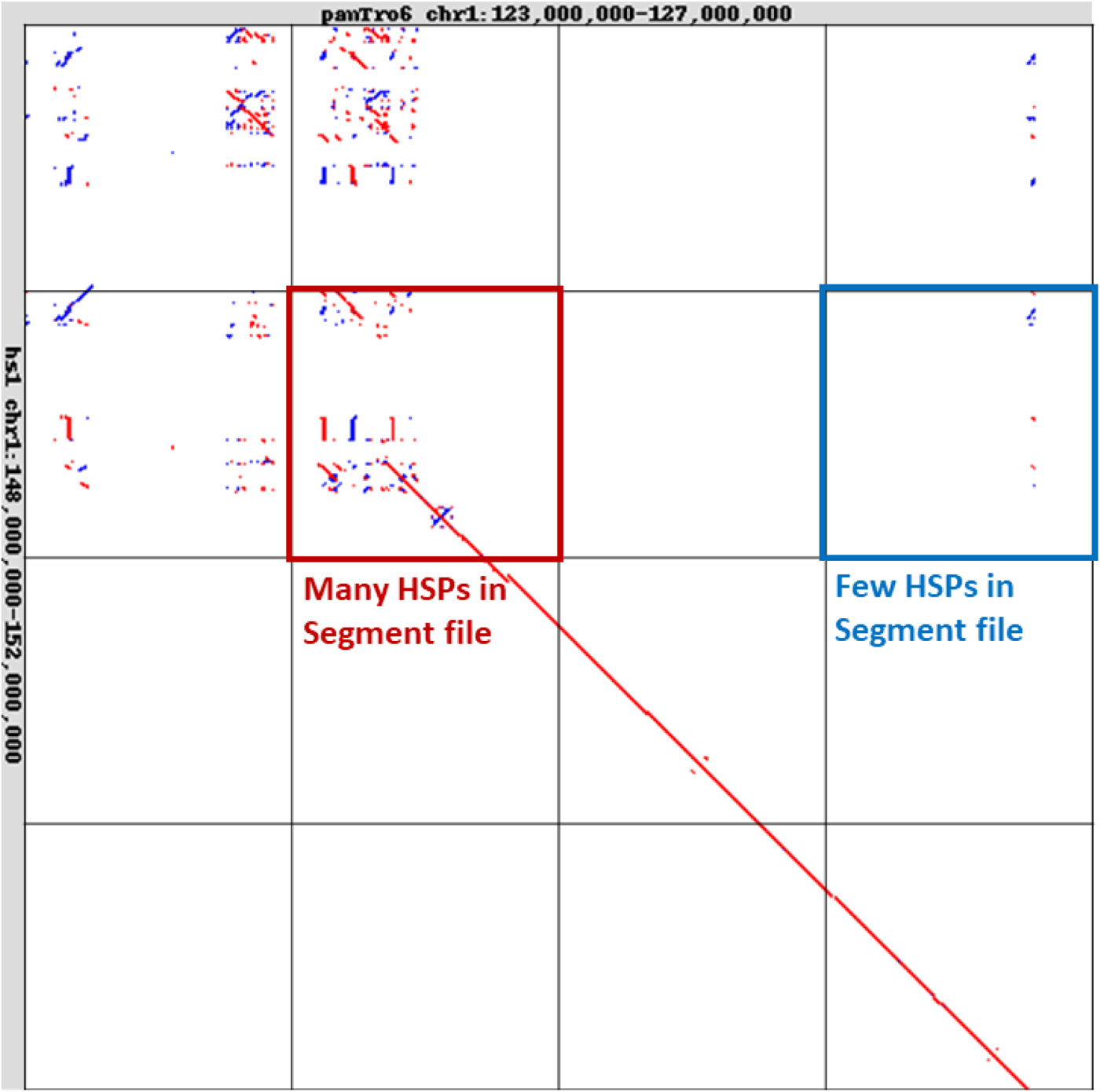
Human – Primate chr1 alignment. Each chromosome has ∼250 million nucleotides. Each segment file contains HSPs from a single chunk. Each box indicates a 10x10Mbp chunk. Some chunks can have very few alignments as shown in the blue box, while others can have significantly more, as shown in the red box.

**Supplemental Figure 2.**
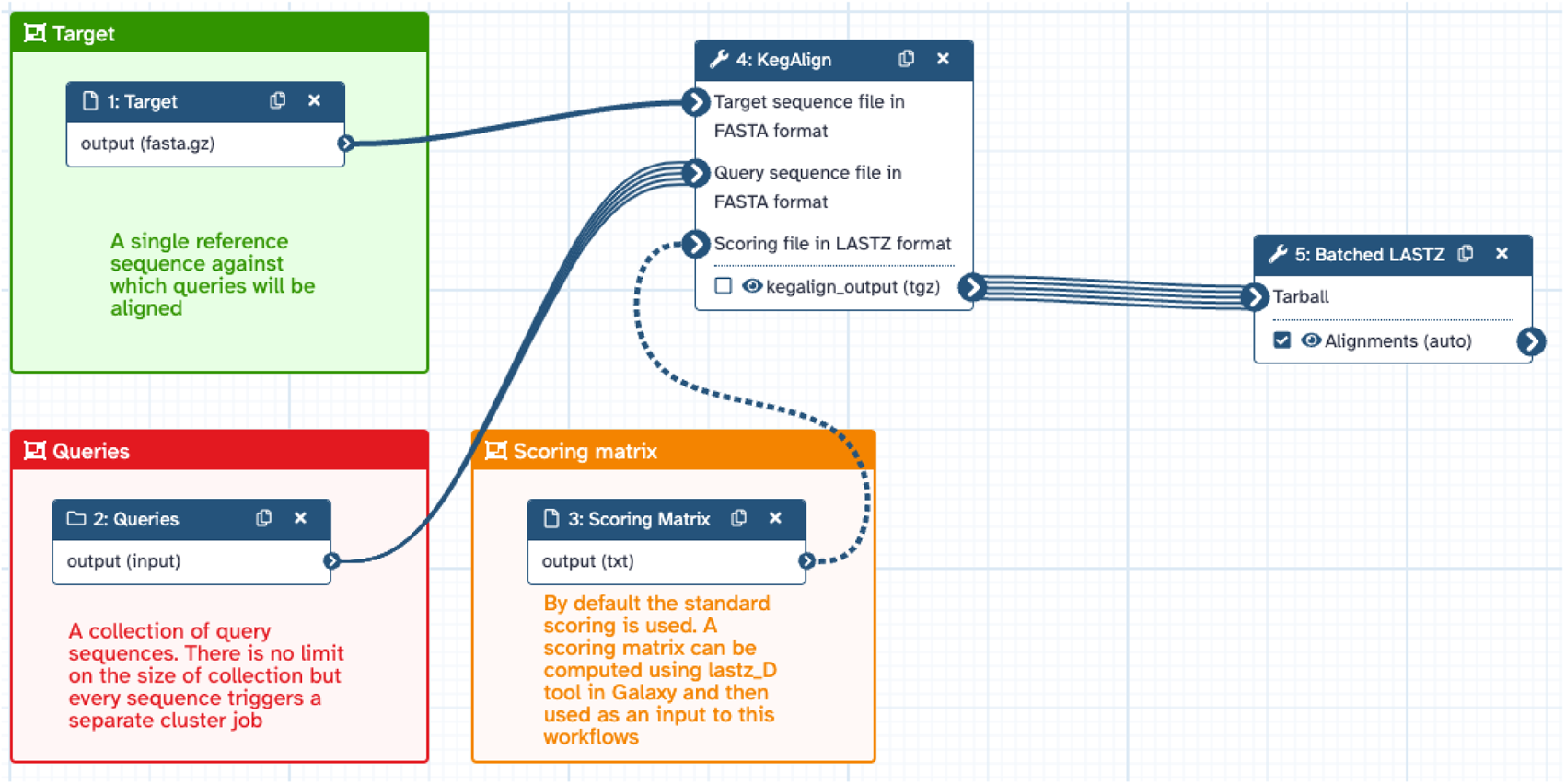
KegAlign workflow available at https://usegalaxy.org/u/cartman/w/cdsoverlaps

